# Atomistic Dynamics of a Viral Infection Process: Release of Membrane Lytic Peptides from a Non-Enveloped Virus

**DOI:** 10.1101/2021.01.07.425769

**Authors:** Asis K. Jana, Eric R. May

## Abstract

Molecular simulations have played an instrumental role in uncovering the structural dynamics and physical properties of virus capsids. In this work we move beyond equilibrium physicochemical characterization of a virus system to study a stage of the infection process which is required for viral proliferation. Despite many biochemical and functional studies, the molecular mechanism of host cell entry by non-enveloped viruses remains largely unresolved. Flock House Virus (FHV) is model system for non-enveloped viruses and is the subject of the current study. FHV infects through the acid-dependent endocytic pathway, where low pH triggers externalization of membrane disrupting (γ) peptides from the capsid interior. Employing all-atom equilibrium and enhanced sampling simulations, the mechanism and energetics of γ peptide liberation and the effect of pH on this process is investigated. Our computations agree with experimental findings and reveal nanoscopic details regarding the pH control mechanism which are not readily accessible in experiments.

## Introduction

Understanding the molecular mechanism by which viruses invade host cells can provide insights for developing anti-viral therapies and drug treatments for diseases caused by these viruses. Enveloped viruses such as human immunodeficiency virus (HIV), Ebola and Influenza breach host cell membranes through a membrane-fusion process catalyzed by fusion proteins in the viral envelope and this process has become increasingly well characterized.(1, 2) However, the mechanisms of cellular membrane penetration by non-enveloped viruses remain enigmatic. Emerging studies suggest that several of these non-enveloped viral families including, *Picornaviridae, Polyomaviridae, Reoviridae, Parvoviridae, Adenoviridae*, and *Nodaviridae* contain a membrane active component of their capsid, which becomes active under endosomal conditions.(3–6)

Flock House Virus (FHV), an insect infecting member of the *Nodaviridae* family serves as model system for investigating the molecular details of cell entry in non-enveloped viruses.(7) FHV can replicate in insects(8), plants(9) and yeast cells(10) and the viral genome consists of two single-stranded positive-sense RNA genomes. The capsid proteins initially assemble into an immature *T=3* icosahedral capsid, composed of 60 identical icosahedral asymmetric units (iASU) and each iASU consist of three chemically identical α subunits, which have 407 residues. The subunits are arranged in three slightly different quasi-equivalent positions (A, B and C). The capsid undergoes maturation to an infectious particle involving autoproteolytic cleavage of the capsid proteins between residues Asn363 and Ala364, catalyzed by Asp75. This results in two particle-associated cleavage products, a mature capsid protein β (residues 1-363) and a 44 residue γ peptide (residues 364-407). The cleavage site (N363/A364) is on the interior of the capsid shell and mutations to either the catalytic D75 residue or the cleaved residue N363, block the maturation cleavage and render the particles non-infectious.(11) The implication is that a dissociated γ peptide is essential for viral infection.

The X-ray crystal structure of the mature capsid was resolved at 2.7 Å resolution and shows that γ peptides are non-covalently associated with the capsid interior (**Figure 1**). While the N-terminal residues (364-385) of γ form an amphipathic α-helix., the predominantly hydrophobic C-terminal region of γ (residues 385-407) are not resolved in the crystal structure, presumably do to a disordered structure and/or conformational heterogeneity across the capsid.(12) Biophysical studies have shown the N-terminal helical region (termed γ_1_) is capable of membrane disruption(13–15) while the C-terminal region of γ plays an important role for specific recognition of the FHV genome and packaging of genomic RNA during assembly.(16) Subsequent studies showed that the full-length peptide has significantly greater membrane lytic activity than γ_1_ and mutations and deletions in the C-terminal region could impair the lytic activity and infectivity. (17, 18) The position of the helices of γ are located in significantly different environments in the three subunits of the iASU due to quasi-equivalence. The A subunits form pentamers and the γ peptides from the A subunits (γ_A_) interact with each other and form a pentameric helical bundle with the helical axes oriented normal to the capsid surface (**Figure 1C**). This organization is stabilized by hydrophilic interactions between peptides and are positioned far from the viral RNA.(19) In contrast, γ peptides from B and C subunits are adjacent to the resolved viral RNA segments and form numerous contacts with the genome.(19, 20) Based on the orientation of the γ_A_ peptides and lack of RNA contacts, it has been speculated that the pentameric helical bundle of γ_A_ peptides are responsible for membrane lysis.(19, 21) This model is supported by studies on Nudaurelia capensis *ω* virus (N*ω*V, *T=4*), where γ peptides are positioned in similar environments (quasi-equivalent).(22) Studies with N*ω*V virus using liposome-based assays show that maximal membrane disruption is acquired with less than half of the subunits of the capsid cleaved,(22) whereas structural studies using time-resolved cryoelectron microscopy (cryo-EM) suggest that subunits surrounding the icosahedral fivefold axis (A subunits) are cleaved much earlier than the subunits positioning in different quasi-equivalent environment of the capsid.(23)

**Figure 1.**
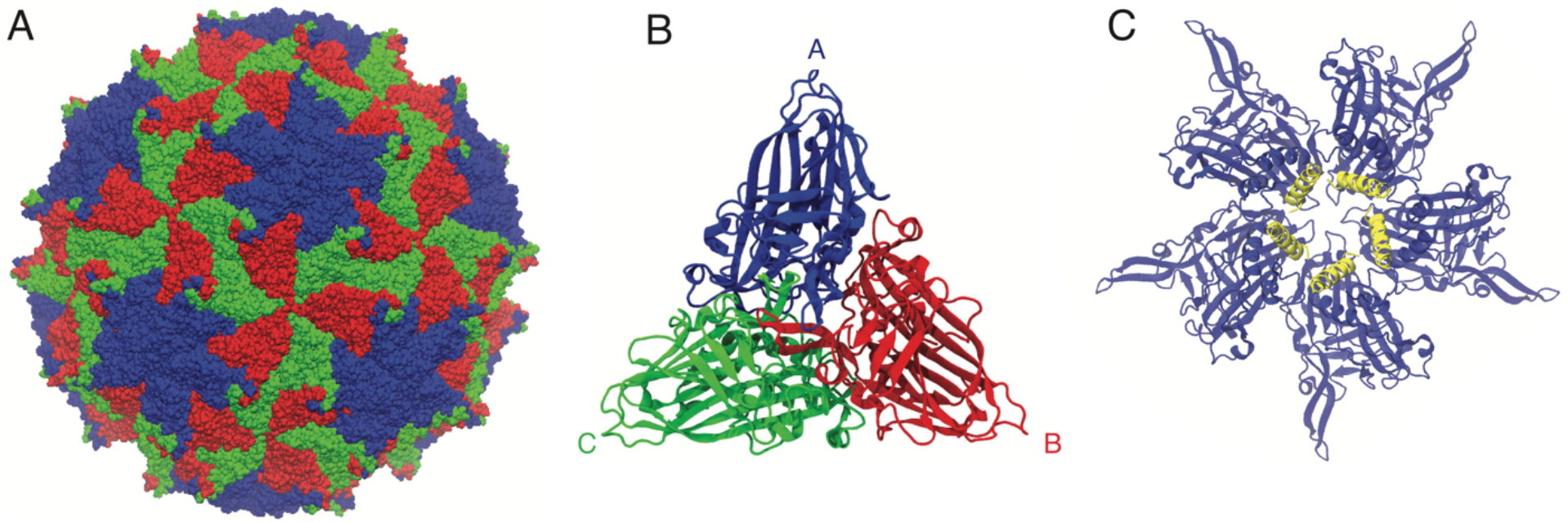
FHV structure. A) *T=3* capsid structure. B) Asymmetric unit. C) Interior view of fivefold vertex. The A-proteins are blue and the yellow helices are the γ peptides. All images are generated from PDBID: 4FTB.

While the FHV capsid crystal structure shows that γ peptides are sequestered inside the capsid,(12) it is believed that the role of the γ peptides in the infection process is to disrupt endosomal membranes and allow the capsid to escape these membrane bound compartments. The proposed model is that low pH induces conformational and/or dynamic changes to the capsid which allow for or promote γ peptides to be externalize from the capsid. This model is supported by several lines of evidence. i) Mass spectrometry combined with limited proteolysis suggest ‘occasional breathing’ or transient exposure of γ peptides on the viral surface.(24) ii) Liposome dye leakage assays showed a pH dependent lytic activity of FHV capsids, with maximal leakage at pH 6.0, consistent with the pH of endosomes.(21) iii) FHV infectivity was also correlated to endosomal pH, as increasing the endosomal pH through the addition of bafilomycin A1, an H^+^

ATPase inhibitor, decreased infectivity.(21) iv) When liposome dye leakage assays were performed at pH 7.0 following preincubation at pH 6.0, for as little as one hour, the dye leakage activity was equivalent to when the leakage assay was performed at pH 6.0.(21) v) bis-ANS, a fluorophore which binds aromatic and hydrophobic residues, had maximal binding at pH 6.0.(21)

While these previous findings provide strong support that FHV has an acid-dependent membrane disruption mechanism involving capsid conformational changes, the nature of structural transition and molecular details are not clearly understood. All-atom molecular dynamics (MD) simulations, sometimes termed a “computational microscope”(25, 26), has emerged as a powerful tool for studying viral systems, as this technique can provide significant insight into the structure and dynamics at a high spatial and temporal resolution, which is typically inaccessible in experiments. Furthermore, MD simulations can go beyond observation of structural features and thermodynamic quantities can be obtained, which can allow a structure-function model to be developed or validated. With ever increasing computational power, several simulation studies have been performed on full virus capsids to investigate the capsid dynamics and physicochemical properties.(27–30) In this present study we use fully atomistic MD simulation to investigate the details of a stage of a viral infection process: the liberation of γ peptides from the interior of a non-enveloped capsid and the effects of pH. Previous simulation studies have examined viral life cycle processes, namely assembly(31–36) and maturation(37–40) using various forms of coarse-grained models and/or implicit solvent models. We believe the current study provides a significant advance in the field by examining a *viral infection process* and its extensive use of atomistic simulations of a large portion of the capsid in explicit solvent through both equilibrium and enhanced sampling simulation methods.

## Results

### pH Effects on the Structure and Equilibrium Dynamics of FHV Capsid

In our study, we begin by analyzing equilibrium simulations of the FHV pentameric capsid system, (**Figure 2A**) at neutral and low pH. These systems were simulated in triplicate for 500 ns each (see **Table 1**). The systems were relatively stable (as measured by C*α* RMSD) during this time scale and similar levels of structural relaxation were observed across the independent trails for each pH, **Figure S1**. The RMSD differences between the neutral and low pH systems are modest, but the low pH system displays a statically significant greater deviation than neutral pH. The averaged RMSD values over the final 300 ns and averaged over the three trails are 0.385 (±0.05) nm and 0.411 (±0.15) nm for neutral and low pH, respectively. We next analyzed the exposure of hydrophobic residues on the capsid surface through SASA calculations. It is experimentally known from bis-ANS fluorescence measurements that low pH induces increased hydrophobic exposure(21), therefore we wanted to observe if our simulations captured this feature. In **Figure 3A** the distributions of solvent accessible surface areas of hydrophobic residues are presented and it is observed that at low pH there is an increase in hydrophobic exposure.

**Table 1.** List of simulation types and lengths.

| Type | pH | Trials/Windows | Simulation Length | Total Simulation Time |
| --- | --- | --- | --- | --- |
| Equilibrium | Neutral | 3 | 500 ns | 1.50 μs |
| Equilibrium | Low | 3 | 500 ns | 1.50 μs |
| SMD - *Single Peptide CV* | Neutral | 20 | 25 - 48 ns | 945 ns |
| SMD - *Single Peptide CV* | Low | 20 | 34 - 47 ns | 889 ns |
| SMD - *Coupled CV* | Neutral | 10 | 27 - 41 ns | 334 ns |
| SMD - *Coupled CV* | Low | 10 | 22 - 34 ns | 284 ns |
| Umbrella Sampling | Neutral | 69 | 100 - 322 ns | 10.67 μs |
| Umbrella Sampling | Low | 70 | 88 - 206 ns | 7.51 μs |
| All Trajectories | | | | 23.63 μs |

**Figure 2.**
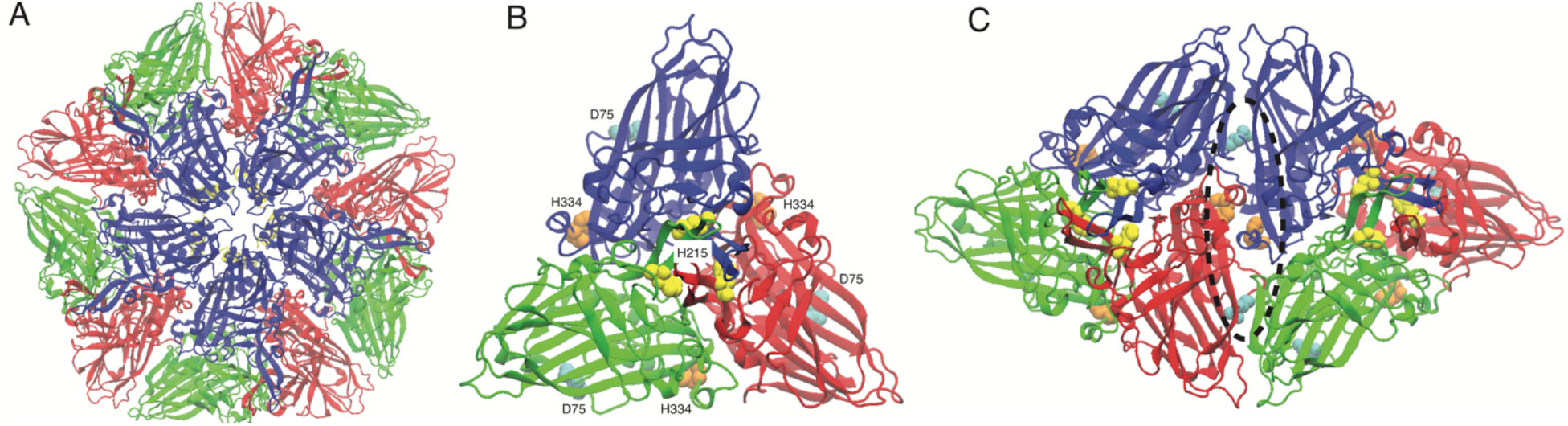
FHV simulation system and titratable residues. A) The simulation system consists of five asymmetric units surrounding a fivefold symmetry axis. The γ peptides associated with the A subunits are retained and can be faintly seen in yellow beneath the fivefold vertex. B-C) Location of residues which change protonation states in low pH model. The residues consist of D75 (cyan), H215 (yellow) and H334 (orange) which are shown in vdW representation. Both a single asymmetric unit (B) and two adjacent asymmetric units (C) are shown to illustrate the location of titrated residues related to neighboring proteins. The H215 residues lie at the center of the asymmetric unit (pseudo threefold symmetric axis), while H334 and D75 lie at the interface between asymmetric units, which is highlighted by the dashed black line. All images are generated in an external to the capsid perspective (looking down on capsid), and are from PDBID: 4FTB.

**Figure 3.**
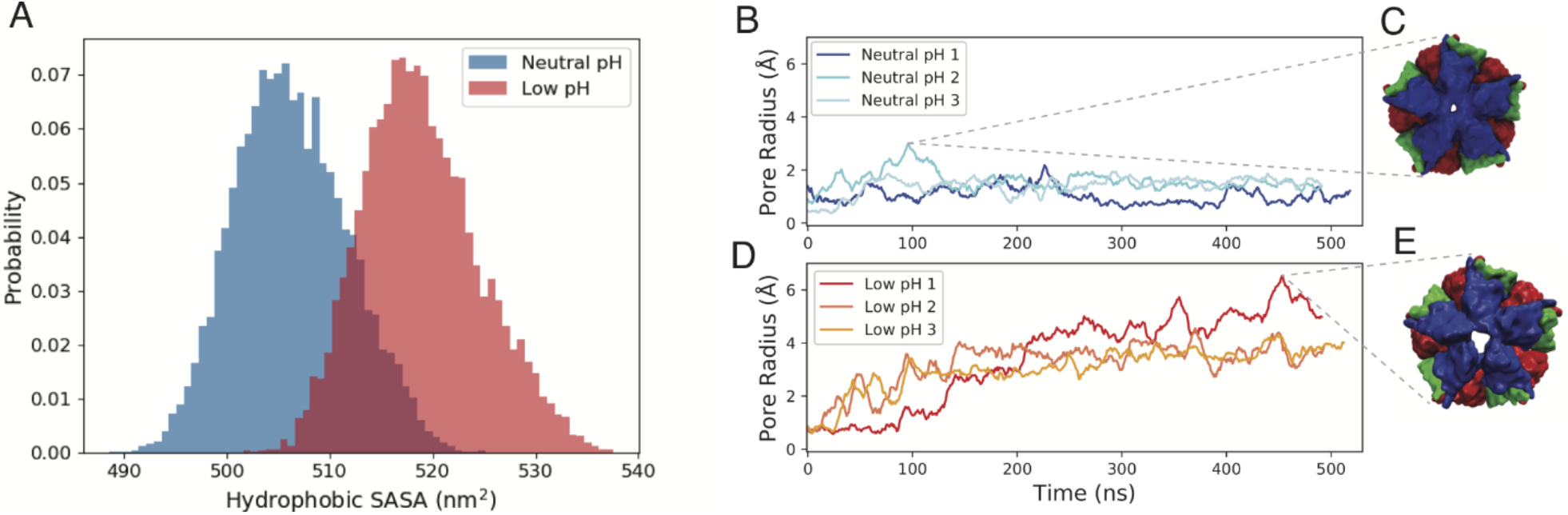
Structural changes induced by low pH. A) The exposure of solvent accessible hydrophobic residues at neutral and low pH. B-E) The pore radius at the fivefold axis is measured for the replicates at both neutral (B) and low pH (D). The structure with the maximum pore radius is shown for neutral (C) and low pH (E) simulations.

While the SASA and RMSD calculations indicate increased conformational changes in the capsid at low pH we wanted to further connect structural changes to potential functional motions of the capsid. Based upon the structural organization of the γ peptides it has been proposed in the literature that γs will externalize through the fivefold symmetry axis,(19, 41) and it has been shown in N*ω*V that pentameric γs are primarily responsible for membrane disruption.(22) Therefore, we examined the dynamics of the capsid at the fivefold symmetry axis to evaluate if low pH induces increased dynamics in the fivefold vertex region. The pore radius at the fivefold vertex was measured during the simulations and the profiles (**Figure 3B**,**D**) show a consistent increase in pore radius at low pH, which is consistent across the multiple trials. The maximum observed pore radius at low pH is 7.3 Å (**Figure 3E**) which is more than double the maximum pore radius at neutral pH (3.5 Å, **Figure 3C**). Furthermore, averaging over the last 300 ns and over the replicates shows the average pore radius at low pH is 4.0 (±0.6) Å, while the neutral pH simulations display an average pore radius of only 1.3 (±0.2) Å. Representative equilibrium trajectories are visualized in **Movie S1** (neutral pH) and **Movie S2** (low pH) to demonstrate the fivefold pore dynamics. These pore measurements reveal a potential γ liberation pathway through the fivefold axis and this pathway is consistent with the previous hypothesis based on the FHV structure-based model.

While we observe structural changes at the fivefold axis, which we believe are functionally significant, the residues which have undergone protonation at low pH are distal to the fivefold vertex (**Figure 2B**). To identify correlations between the titratable residues and capsid functional motions, we examined the structural dynamics of the systems by calculating secondary structural (SS) changes and linear mutual information. In **Figure 4A** we present the change in secondary structure of the A subunits relative to the neutral pH capsid, collected over the last 300 ns of the simulation trajectories. In **Figure 4B-C**, we show the location of residues with significant (> 10 %) changes in SS. The residues which display significant changes in α-helix or β-sheet content are 91-97, 169-171, 178, 203-204, 210-211, 244, 328-332, 334-335, 352-353 and 357-358. We see that these residues are proximal to the two histidine residues (H215 and H334) and participate in the protein-protein interfacial contacts. These subtle conformational changes could have a profound effect on the collective dynamics of the capsid. Furthermore, it appears there is a connection of residues, emanating from H334, which display differential SS changes and propagating toward the fivefold vertex (**Figure 4C**). Interestingly, we did not find any change in SS in the neighboring regions of the other protonated residue, D75.

**Figure 4.**
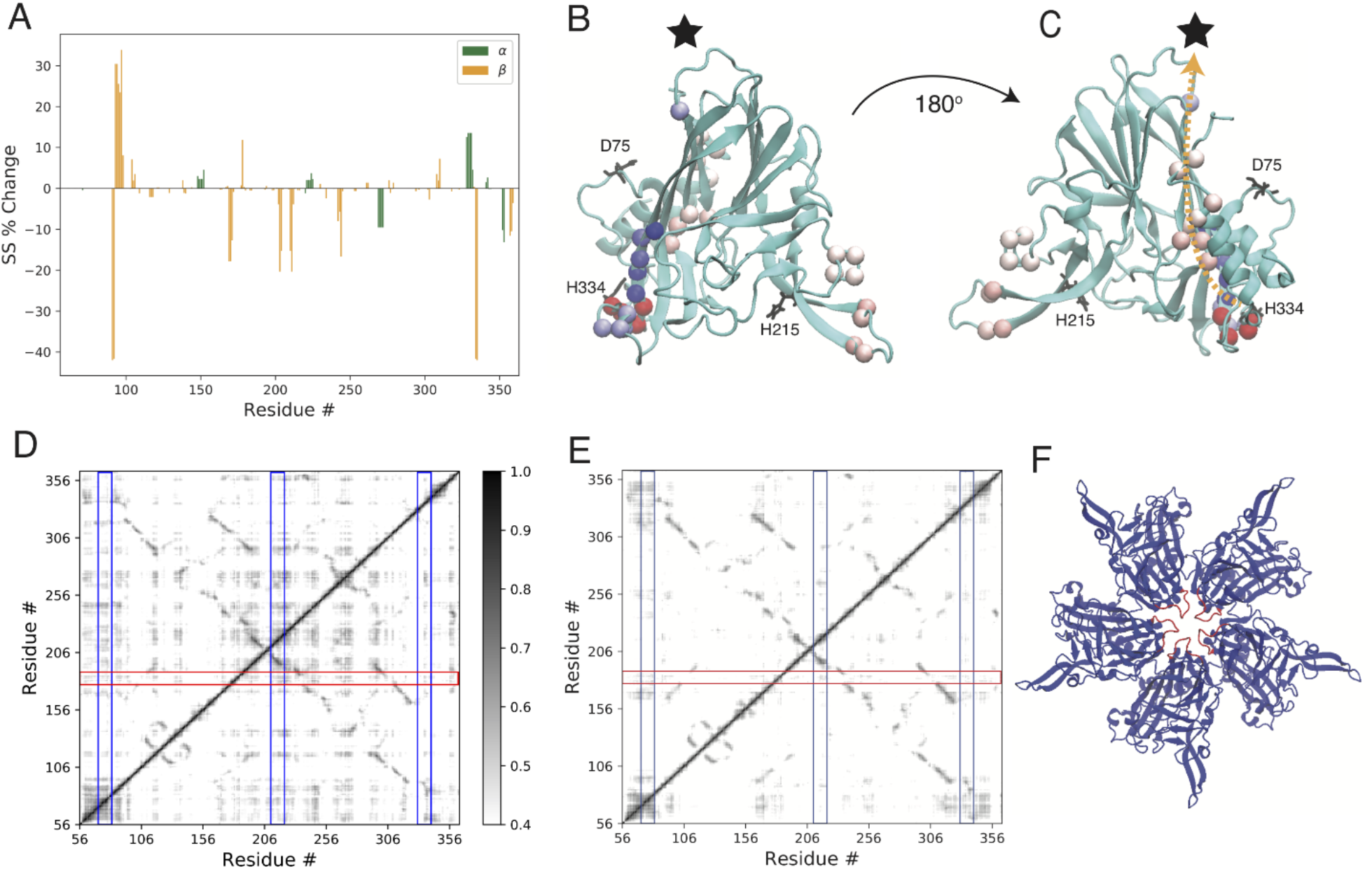
pH effects and dynamic couplings. A) The change in secondary structure (SS) in the A subunit in going from neutral to low pH. B-C) The C*α* atoms of residues with SS changes greater than 10% are displayed as vdW spheres on the A subunit structure. Colors scale is from red (largest decrease) to blue (largest increase). Top down (B) and internal (C) views are presented. The black star indicates the location of the fivefold vertex, and the orange dashed line is drawn to illustrate a pathway from H334 to the fivefold vertex. Residues which undergo protonation changes are labeled. D-E) The linear mutual information at neutral (D) and low pH (E). The correlations with less than 0.4 are set to zero to highlight stronger correlations. The blue vertical boxed regions are surrounding the titrating residues (D74, H215 and H334) and the red horizontal box is around residues 178-186, which is a loop at the fivefold vertex. F) The boxed red region in (D-E) corresponds to the red loop shown on the pentameric structure.

We performed linear mutual information (LMI) analysis to characterize the correlations inside the capsid and the corresponding matrices are shown in **Figure 4D-E**(42, 43). In general, more mutual information is observed across many different regions of capsid at neutral pH. Interestingly, there is significant mutual information between the inner most (“gate-keeper”) loop of the fivefold axis with different regions of the capsid, but these correlations are diminished at low pH. The region around H215 also show strong correlations across the capsid, including with the gate-keeper loop and again most of those correlations are diminished at low pH. H215 is located at the center of the asymmetric unit (pseudo threefold axis, **Figure 2B**) and may serve as a hub residue capable of modulating protein-protein dynamics throughout in the capsid. Our results lead us to hypothesize that under low pH conditions the capsid motion of the fivefold axis is not concerted with the other regions of the capsid, which we suspect may lead to pore formation at the fivefold vertex. Our interpretation of the LMI and SS analyses is that the capsid is more connected and possibly more rigid at neutral pH, and at low pH the protonation of the histidine residues leads to a disruption of the network. In particular these results point toward a model in which under low pH conditions the capsid motion of the fivefold axis is not concerted with the other regions of the capsid, which we suspect leads to weaker/softer interactions at the fivefold vertex that is supported by the pore analysis (**Figure 3B-E)** and further investigated in the following sections.

### γ liberation pathways from Non-Equilibrium Simulations

To gain insight into the molecular details of γ liberation, we performed biased (steered MD) non-equilibrium simulations to generate translocation of the γ peptides from the interior (under) of the capsid to the exterior (above). We performed the biasing in two separate approaches. In one approach (referred to as *coupled CV*) the collective COM of the five peptides was biased; in the second approach (*single peptide CV)*, the COM of a single peptide was biased. A representative trial for each biasing method is presented in **Movie S3** (*coupled CV*) and **Movie S4** (*single peptide CV*) for the low pH system. The *coupled CV* was partially motivated by speculation in the literature that the five peptides would move as a unit/bundle and externalize simultaneously(19, 41). Using the *coupled CV* was an attempt to evaluate the bundle hypothesis, where if there interpeptide interactions were sufficiently strong the peptides they would move collectively (stay as a bundle) and not separate. We conducted 10 independent pulling simulations for each system (neutral and low pH capsid) and all simulation trajectories clearly show that γ peptides were not liberated simultaneously. This finding contradicts the previous bundle hypothesis. We have calculated the nonequilibrium work done by the pulling force required for liberation of the first γ peptide using the *coupled CV* (**Figure S2A-B**). We observe that the low pH environment reduces the work required for externalization in a statistically significant manner (**Figure S2B**). We also examined the maximum force observed during the SMD trials but did not find the pH significantly affected this quantity (**Figure S2C**). For the trials at each pH which displayed the lowest work values, we continued the externalization until all five peptides were externalized. For this calculation, we removed the γ peptides from the system once they were liberated from the capsid and then restarted the SMD run with the *coupled CV* based on the remaining peptides COM. The work required to liberate all five peptides is reduced in low pH compared to neutral pH (**Figure S3**). Interestingly, the largest amount of work is required to release the first peptide and then the amount of work subsequently decreases as the following peptides are liberated. This results indicates the release of the first peptide is critical and possibly the rate-limiting step to achieve membrane disruption, which motivates further investigation into the mechanism and energetics of the release of a single peptide.

To further explore the γ liberation pathway, we performed 20 independent steered MD simulations at each pH, using the *single peptide CV*. The temporal evolution of external work done by the pulling force for the individual trajectories is shown in **Figure 5A**, and a clear separation (non-overlapping distribution) is observed. The final work values are compared in **Figure 5B** and at neutral pH the average work is 403 (± 8) kcal/mol and at low pH the average is 291 (± 5) kcal/mol, a reduction of over 100 kcal/mol when the pH is lowered. The SMD bias velocity was 0.1 nm/ns in both biasing approaches (*coupled* and *single peptide* CVs). To check that the work trends are not highly sensitive to the pulling velocity a series of SMD trials were performed at different pulling rates (0.2, 0.4, 0.8 nm/ns, see **Figure S4**), which demonstrate that the results are not unique to the pulling velocity chosen. The average maximum force is significantly higher (1187 ± 21 pN) at neutral pH compared to low pH (948 ± 35 pN), indicating that stronger (or more) interactions resist the γ liberation process in neutral conditions(**Figure 5C**). The maximum forces are large, however other viruses, including *T=3* non-enveloped capsids have been shown to be able to withstand applied forces in the range of 1000 pN without undergoing irreversible deformation. This is based on atomic force microscopy (AFM) nanoindentation experiments and CG MD simulations.(44–47) The SMD results appear consistent with the linear mutual information analyses (**Figure 4D-E**), which indicated the system was more strongly correlated and rigid at neutral pH. Hence, our multiple SMD simulations confirm that the energetic requirements to release a γ peptide are significantly reduced under low pH conditions which is in qualitative agreement with the previous experimental work.(21) While the non-equilibrium work values can be related to the equilibrium free energy changes through the Jarzynski equality,(48) this method is difficult to converge and can require greater than 100 trials.(49, 50) Therefore we next attempted to estimate the equilibrium free energy profiles of γ liberation by performing extensive umbrella sampling simulations along the minimum work pathways under the different pH conditions.

**Figure 5.**
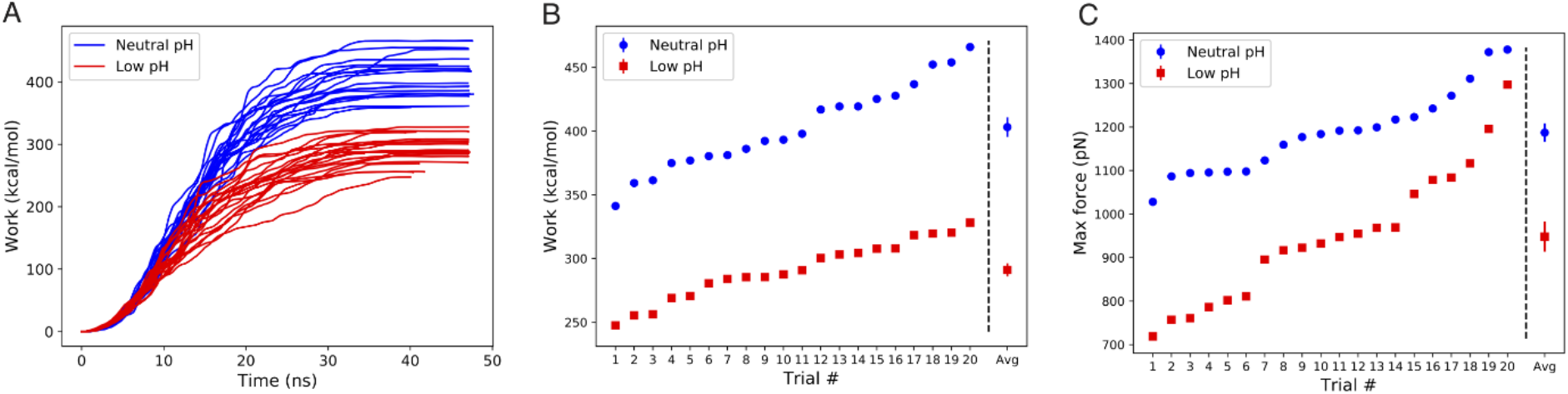
SMD results using *single peptide CV*. A) The work versus time for the 20 trials at each pH. B) The final work values are sorted and presented from smallest to largest. The average over the trials is shown at the far right. C) The maximum force values are presented and ordered from smallest to largest. The average over the trials is shown at the far right. In (B-C) the error bars on the average values are the standard error of the means.

### pH Effects on energetic barriers for γ liberation

To calculate a free energy profile of γ liberation under the different pH conditions, the lowest work trials at each pH (**Figure 5A**) were selected as initial pathways. These pathways were then subjected to extensive umbrella sampling simulations to allow for the systems to relax from the non-equilibrium pathways to a locally equilibrated pathway. To detect that each window had properly equilibrated and was well sampled, autocorrelation based timeseries analysis was performed.(51) Umbrella windows were run for up to 322 ns in order to retain approximately 20% (18.9% was the minimum) of the data set and have a sufficient (> 35) *N*_*eff*_ in each window (**Figure 6A-B**). A wide range of equilibration times are observed, but we consistently observe fast equilibration times in the high numbered windows (>60), where the peptide is externalized and dissociated from the capsid. Since the peptide is effectively in solution at this stage of the pathway, it is reasonable to expect these simulations would rapidly equilibrate.

**Figure 6.**
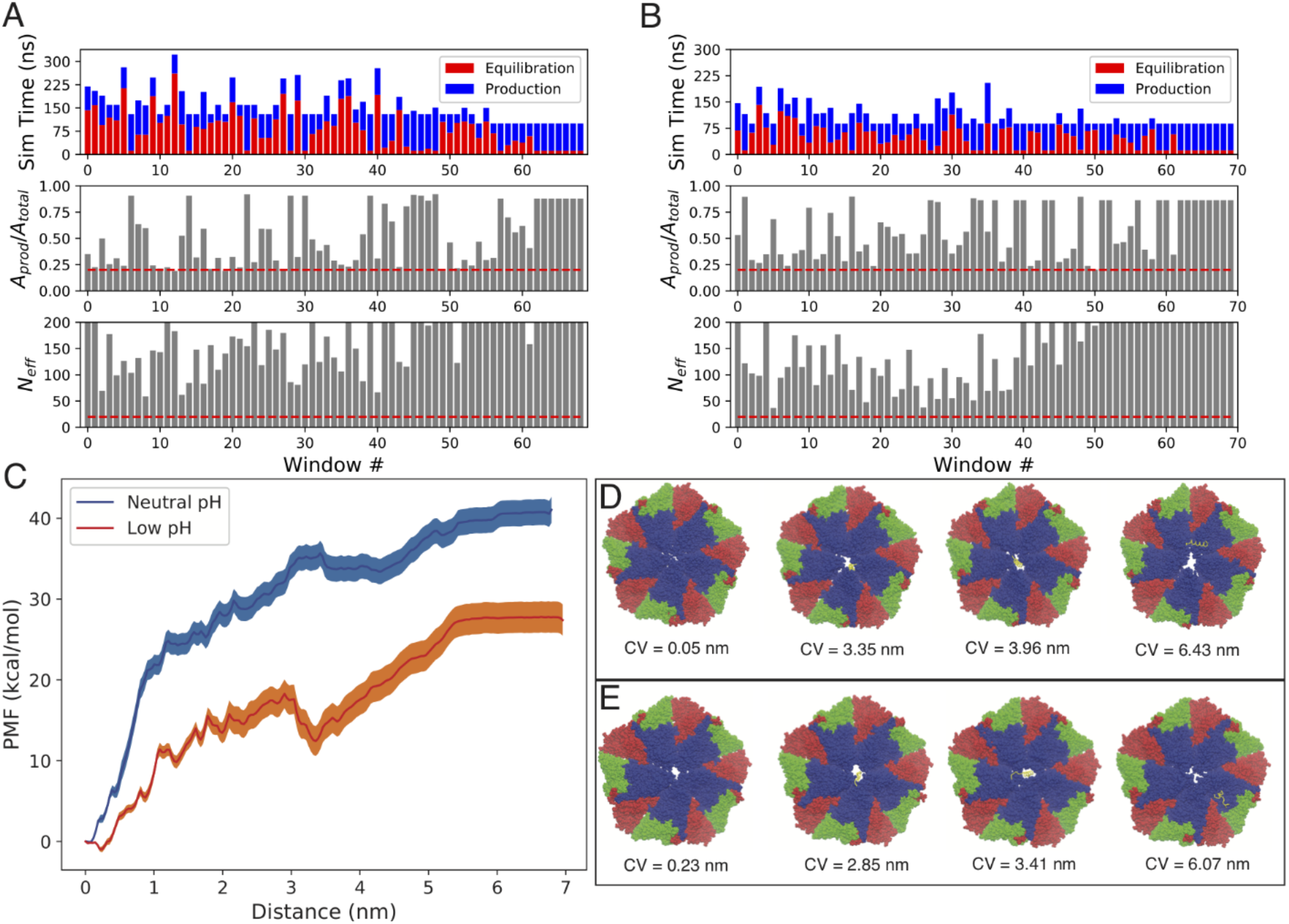
Energetic Analysis of γ liberation pathways. Time series analysis of umbrella sampling data at neutral (A) and low pH (B). The top plot shows the breakdown between equilibration and production data for each window. The middle panel shows the ratio of production to total simulation data, with dashed red line drawn at 20%. The bottom panel shows *N*_*eff*_ with dashed red line drawn at *N*_*eff*_ = 20. The y-axis is terminated at 200, to highlight lower sampled windows, though many windows have *N*_*eff*_ > 200. C) PMF force at neutral and low pH. Shaded region represents the standard deviation obtain from error analysis using1000 bootstrapped profiles. D-E) Structures along the γ liberation pathway at neutral pH (D) and low pH (E) pathways. Snapshots are the final frame from a given umbrella window.

The free energy profiles were computed from the equilibrated portion of data in each window, which totaled 5.0 μs at neutral pH (window minimum sampling time = 25 ns) and 4.2 μs at low pH (window minimum sampling time = 18 ns). The resultant profiles (**Figure 6C**) show a significant reduction in the barrier (Δ*G*_*barr*_) and overall (Δ*G*_*ext*_) free energy change when the pH is reduced. At neutral pH a sharply increasing profile is observed in the first 1 nm, followed by a more rugged and more gradual increase until reaching a barrier of ~36 kcal/mol at ~3.4 nm. This is followed by a slight decrease in the PMF reaching a local minimum of ~34 kcal/mol at ~4.0 nm. The PMF then increases reaching ~40 kcal/mol when the peptide fully dissociates from the capsid around 6.0 nm. Whereas in low pH a more gradual and jagged profile is observed reaching a barrier of ~18 kcal/mol at 2.9 nm. This barrier is followed by a sharp decrease in the PMF to ~12 kcal/mol, where the system reaches a metastable state at ~3.4 nm. The metastable state is shown in **Figure S5** and is characterized by partial exposure of the γ peptide onto the capsid exterior surface. This is followed by a gradual increase in energy to ~28 kcal/mol at the fully dissociated state at ~6.0 nm. Snapshots corresponding to the initial, barrier, metastable and final states are shown in **Figure 6D-E** for both systems. Overall, we observe Δ*G*_*barr*_ is reduced by half (36 kcal/mol at neutral, 18 kcal/mol at low pH) and Δ*G*_*ext*_ is reduced from 40 kcal/mol to 28 kcal/mol when the system pH is reduced.

## Discussion

Several non-enveloped viruses including *Rhinovirus, Hepatitis A, Reovirus, Parvovirus, Adenovirus* and FHV contain membrane disrupting components of their capsids which are triggered to release/externalize under low pH (endosomal) conditions.(4, 5) However, the molecular mechanisms that underlie this process remain unclear and are challenging to address using current experimental techniques. The rapid progress in computational power and simulation algorithms now permits MD to investigate these complex biomolecular processes and can provide important insights at atomic resolution into the mechanism of action, where experimental characterization is rather difficult if not impossible.(52)

In the present study, we have investigated the mechanism of lytic peptides liberation from the FHV capsid and the role of pH in affecting the structural dynamics and energetics of the liberation process through a multiscale atomistic simulation approach including equilibrium simulations, steered MD and extensive umbrella sampling. Based on the FHV structure-based model, it is hypothesized that γ peptides located under the fivefold axis are liberated from the capsid to disrupt cellular membranes and permit infection. It is known from *in-vitro* studies that γ-peptides disrupt membranes(13–15) and do so with maximal efficiency at pH 6.(21) Furthermore, when the γ peptides are not cleaved from the capsid protein the particles are effectively non-infectious,(11) substantiating the critical role these peptides play in viral infection. Interestingly, from our 0.5 μs long equilibrium simulations we are able to observe pore opening at the fivefold axis under low pH conditions (**Figure 3, Movie S2**). Our analysis demonstrates that at low pH, protonation of histidine residues affect (decrease) allosteric communications across the capsid, which may lead to the gate-keeper loop opening (pore formation) in the capsid (**Figure 4**). The role of the two histidines as “pH sensor” residues which can affect capsid dynamics is supported by conservation of the histidine positions in closely related *nodaviruses* (see multiple sequence alignment in **Figure S6**). Though the formation of a pore in the low pH capsid is an exciting observation, we do not observe spontaneously translocation of the γ peptides, which would presumably require simulation time scales beyond our current computational capabilities.

It is therefore informative to understand how pore formation in the low pH capsid affect the energetics of γ liberation process. In this study the γ liberation process is characterized through a series of SMD runs, followed by extensive umbrella sampling along the most favorable pathways. Similar approaches combining SMD with umbrella sampling have been successfully applied to study processes of other large biological complexes such as the chromatophore(53) and membrane transporters.(54) The SMD simulation results provide a qualitative but conclusive demonstration that the work necessary for externalizing γ peptides is distinctly lower at pH 6 than at pH 7, which is well corelated with the optimal pH of FHV membrane lytic activity.(21) In our umbrella sampling approach we allow for long relaxation times (> 100 ns) and select only well equilibrated data, based upon autocorrelation analysis, to reconstruct the free energy profiles. We observe significant reductions in the barriers and overall free energy change required to externalize a peptide at low pH.

While the overall free energy changes we observe, even at low pH, are positive there are a few factors which could contribute to this result. One is that the initial pathways based upon SMD are far from equilibrium. Even though the umbrella sampling simulations are quite long, compared to previous all-atom studies on capsids, (37, 55) there could be residual strains in the system which are not fully relaxed leading to overestimation of the free energy profiles along the reaction coordinate. Another, more physical consideration, is to contemplate the closed system of the capsid within a membrane bound vesicle/endosome. The process of liberating the peptide would involve the free energy cost of externalizing the peptide, but also the free energy of partitioning the peptide into the membrane, which is known to be favorable. Experimental measurements indicate the Δ*G*_*part*_ for γ_1_ partitioning into lipid bilayers to be approximately −8 kcal/mol(14), while CG simulations have estimated the ΔG of γ_1_ binding to different bilayers in the range of −15 kcal/mol to −22.5 kcal/mol, when the peptide is in a helical state.(56) This computationally computed free energy of binding is comparable to the computed free energy of externalization (28 kcal/mol) at low pH. Furthermore, the peptide is partially exposed in the metastable state (**Figure S5**) and if the virus were to closely approach the membrane the peptide might be able to initiate contact with the bilayer before fully dissociating from the capsid. In this scenario the overall process could be favorable as the externalization free energy cost to reach the metastable state is only 12 kcal/mol at low pH but is far too high (34 kcal/mol) to be favorable at neutral pH. It remains unclear if the peptides fully dissociate from the FHV capsid or if they are remaining in contact with the capsid. There would be a functional benefit for γs to remain in contact with the capsid, as this would allow for the capsid to be proximal to the location of membrane disruption. In poliovirus there is evidence to support membrane disrupting peptides are involved in umbilical connections between the capsid and bilayer.(57)

In this work we have explicitly modeled a curvature restrained fifteen subunit subsection of the FHV capsid, which represents 1/12 of the complete capsid. It is known that large biomolecular assemblies, such as virus capsid exhibit long range collective motions, which can often be low frequency (low energy) motions that can couple to functional dynamics of the systems.(27, 38, 58) It is possible in the context of the complete capsid that collective motions would facilitate the peptide release, which are damped in our reduced system, however our LMI analysis indicates that the capsid becomes less collective at low pH (**Figure 4**). Another component to the complete virus, which is neglected in the current study, is the viral genome. While we cannot discount the possibility that low pH induces conformational changes in the genome, which then influence the pathway and energetics of γ liberation, there is evidence that γ release and genome release for FHV are separate events. This is based upon a recent study that used heat to trigger genome release from the FHV capsid.(59) During the heating they observe multiple transitions measured by differential scanning calorimetry (DSC) for the wild-type virus, but only a single transition for maturation defective mutants. The first transition for the wild-type particle corresponds to γ release as detected by anti-γ immunolabeling and is independent from the genome exposure transition.

There is increasing interest in understanding asymmetric dynamical processes in highly symmetric virus structures as these are functionally relevant processes and now both MD and cryo-EM are capable of characterizing these phenomena.(60, 61) The present study provides the first free energy profile for membrane lytic peptide liberation from a nonenveloped virus at different pH values. The results provide strong evidence that lowering the pH induces increased fluctuations at the capsid fivefold vertex and reduces the energy requirement to externalize the lytic peptide. We identify a metastable state at low pH which positions the peptide in an externalized but capsid associated state. When the membrane binding step is taken into consideration the overall process of transitioning a peptide from the capsid interior to the membrane may approach a negative ΔG, especially if the membrane binding were to from the metastable state. Overall, our simulation study results qualitatively agrees with the previous experimental observation and elucidates several molecular-level aspects of how low pH can trigger capsid structural rearrangement, facilitating externalization of membrane lytic peptides that can disrupt the endosomal membrane and allow for viral infection to proliferate.

## Methods

### System Preparation

In this study we modeled a subsection of the complete FHV capsid, comprising five asymmetric units surrounding a fivefold symmetry axis (**Figure 2A**). The initial coordinates used were taken from the 2.7 Å resolution X-ray crystal structure (PDB ID: 4FTB). The N-terminal tail of the A, B and C subunit, and C-terminal regions of the γ peptides are unstructured and not resolved in the crystal structure. The unstructured regions of the capsid, residues1-55 of chain A, residues 1-52 of chain B, residues 1-54 of chain C and residues 385-407 of the γ peptides were not modelled into this structure to avoid artifacts from generating initial conformations of large undetermined regions. Four Ca^2+^ ions are bound to each iASU, and these ions were included in our starting structure. To maintain the capsid curvature and mimic a full-capsid model, weak positional restraints (force constant of 500 kJ mol^−1^ nm^−2^) were applied to several C_α_ atoms, positioned along the edge of the pentameric structure. Restrained atoms are listed and shown in **Figure S7**. Low pH conditions were modeled by protonating residues D75, H215 and H334. The low pH system is attempting to model pH~6 which is the environment where FHV capsids have maximal lytic activity.(21) There are only two histidine residues in each subunit and the choice to protonate the histidines is based upon assuming a p*K*_a_~6.5.(62) D75 was protonated based upon a previous p*K*_a_ prediction of ~6.0 and the proposed reaction mechanism for γ cleavage involves a protonated D75 acting as a hydrogen donor.(63) Each system was solvated with the TIP3P water model(64) and was neutralized by the addition of Na^+^ ions and additional Na^+^ and Cl^−^ ions were added to generate a physiological ion concentration of 0.15 M NaCl. The total system sizes were approximately 390,000 atoms.

It is worthwhile to discuss the use of fixed protonation states in modeling different pH environments as we done in this work. Sophisticated methods have been developed to model constant pH environments in atomistic MD simulations, where residue titration states are coupled to the system dynamics. (65, 66) This approach has been applied to the study of virus capsid systems, (37, 67) however those studies were performed on a single asymmetric unit using rotational symmetry boundary conditions and in an implicit solvent environment. The constant-pH MD method has been recently extended to be compatible with explicit solvent either through a hybrid implicit/explicit solvent approach(68) or through the multi-site λ dynamics formalism.(69) While the explicit solvent constant pH approaches are very promising, especially for membrane-protein systems, they still come with a high computational demand. To our knowledge the application of these methods have been limited to system sizes in the range of 50,000 atoms and with individual trajectory lengths no longer than 62 ns.(70–73) In our study we are able to achieve simulations times up to 500 ns and total sampling for over 23 μs for an explicitly solvated system of nearly 400,000 atoms, which enables us to uncover principals that underlie acid-induced externalization of γ peptides from the capsid and qualitatively agree with previous experimental observations.

### General Simulation Protocol

All simulations in this study were performed with the GROMACS 2018 simulation package(74) using the CHARMM36 all-atom force field.(75) All simulation types, lengths and trials are listed in Table 1. The systems were first energy minimized using the steepest-descent algorithm followed by 100 ps of NVT (300 K) followed by 100 ps of NPT (300 K, 1 atm) equilibrations. In the NVT and NPT equilibrations, the nonhydrogen (heavy) atoms of the protein were restrained with a force constant of 1,000 kJ mol^−1^ nm^−2^. Production MD simulations were performed for 500 ns at a temperature of 300 K and a pressure of 1 atm. Constant temperature was maintained using the Nose-Hoover thermostat(76, 77) with a coupling time constant of 1 ps and constant pressure was maintained using the Parrinello-Rahman barostat(78) with a coupling constant of 5 ps. Simulation timestep was 2 fs and waters were kept rigid using the SETTLE algorithm.(79) Non-water bonds involving hydrogen atoms were held fixed using the LINCS algorithm.(80) Electrostatic interactions were calculated by using the particle-mesh Ewald (PME) method(81) with a real-space cutoff of 12 Å. The cutoff for Van der Waal interactions was set to 12 Å, with smoothing starting at 10.5 Å.

### Steered Molecular Dynamics (SMD) Simulation

Externalization of γ peptides from the capsid were characterized through a series of steered MD (SMD) simulations, where a constant velocity pulling restraint was applied to a collective variable (CV), which was the center of mass (COM) of the backbone C_α_ atoms of the γ peptides to produce movement along the z axis. The biasing potential (*U*_*SMD*_) for SMD is:

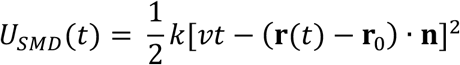

where, *k* is the pulling force constant and *v* is the pulling rate. **r**(*t*) is the position of the CV at time *t* and ***r***_0_ is the initial value of the CV, **n** is the direction of pulling, which in this study was set as the unit vector in the Z-direction. Starting conformations for SMD simulations were randomly chosen from the last 100 ns of the equilibration simulations. The majority of simulations used a pulling velocity of 0.1 nm/ns and the pulling force constant was 1,000 kJ mol^−1^ nm^−2^. Two different CVs were examined, CV_1_ (*coupled*), in which the COM position of all five peptides was collectively biased and CV_2_ (*single peptide*) where a single peptide was chosen and it’s COM position was biased. The non-equilibrium work (*W(t)*) is calculated by integrating the time-dependent force (*F(t)*) acting on the biased CV,

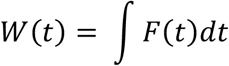

using the cumtrapz function in MATLAB. It is worthwhile to mention that the non-equilibrium work profiles derived from SMD simulations are dominated by the dissipative energy component which is stochastic in nature. Therefore, we performed ~40 SMD simulations for each system using different CVs, different starting conformations and different pulling velocities (0.1-0.8 nm/ns, **Figure S4**) to enable reliable observations of these systems.

### Umbrella Sampling Simulation

We have performed umbrella sampling simulations(82–84) using GROMACS 2018, to estimate the free energy change associated with γ liberation from the capsid at neutral and low pH. A harmonic umbrella potential (force constant of 2092 kJ mol^−1^ nm^−2^) was used to restrain the COM of a single liberating γ peptide along z axis. Umbrella windows were spaced 1 Å apart and spanned the entire γ liberation pathway (~ 7 nm), resulting in 69 windows for the neutral system and 70 windows for the low pH system. Each umbrella window was sampled for at least 88 ns, amounting to a total of 18 μs simulation time. To improve the estimate of the potential of mean force, (PMF) we have extracted only equilibrated samples from each window using the timeseries module of the pymbar package.(51) We have run each umbrella window until approximately 20% of the data is selected as equilibrium samples and there are at least 35 effectively uncorrelated samples (*N*_*eff*_) for each window. *N*_*eff*_ is defined as

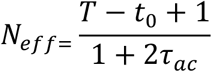

where *T* is the total simulation length, *t*_*0*_ is the equilibration time and *τ*_ac_ the decorrelation time of the biased CV.(51) A minimum equilibration time of 12 ns was applied to windows with *t*_*0*_ less than 12 ns. The unbiased free energy surface was constructed from the umbrella histograms, using the weighted histogram analysis method (WHAM)(85) implemented in GROMACS gmx wham tool(86). All umbrella histograms were visually inspected to ensure significant overlap between the histograms from adjacent simulation windows. The PMFs were converged at 10^−6^ kcal mol^−1^ tolerance. We have estimated the statistical uncertainty of our free energy profile using Bayesian bootstrap of complete histograms method, freely distributed with the GROMACS gmx wham tool(86) and 1000 bootstraps were generated to calculate the statistical error.

### Trajectory Analysis

Root-mean-square-deviation (RMSD) of the capsid C_α_ atoms was calculated with respect to its initial state using the GROMACS tool gmx rms. Translational and rotational superposition of the capsid with respect to its initial state is performed, prior to RMSD calculation. Hydrophobic solvent accessible surface area (SASA) was calculated using the VMD package(87) by using a spherical probe of 1.8 Å. The definition of hydrophobic residues for this calculation is Ala, Leu, Val, Ile, Pro, Phe, Met and Trp. The calculation was performed over the A, B and C subunits but did not include the γ peptides. The final 300 ns of each of the equilibrium simulations at each pH (900 ns total at each pH) was analyzed. Analysis of the pore radius at the fivefold vertex was performed on the capsid over the multiple equilibrium simulation trajectories using the program HOLE(88), implemented in MDAnalysis(89). This pore calculation was performed every 1 ns over the course of the multiple 500 ns equilibrium trajectories at each pH. Residue-wise secondary structural propensity of the capsid subunit A protein was computed using the Dictionary of Secondary Structure in Proteins (DSSP) implemented in GROMACS do_dssp tool.

Mutual Information quantifies the amount of information in one unknown variable (*X*_*j*_) obtained from the state of another known variable (*X*_*i*_). We have calculated the linear contribution (LMI) *(I*_*lin*_*(X*_*i*_,*X*_*j*_)) using bio3d(42, 43) and based on the following formula:

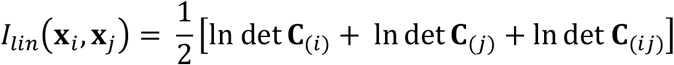

where 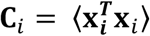 is the marginal covariance matrix, **C**_*ij*_ =: ⟨**x**_*i*_**x**_*j*_⟩^3^⟨**x**_*i*_**x**_*j*_⟩= is the pair covariance matrix and **x**^*i*^ is the positional fluctuation of the C*α* atom of residue *i*. Unlike, general mutual information analysis, this method does not require a computationally expensive entropy calculation and provides a lower bound on the mutual information estimate. For both the secondary structure and the LMI analyses the data was averaged over 4.5 μs of data at each pH. This amount of data arises from using the last 300 ns of each of the three trials and averaging over the five A subunits in our simulation system.

## ACKNOWLEDGMENTS

We would like to thank Kevin Boyd for providing guidance and scripts for the timeseries analysis. We also would like to acknowledge Allyn Brice, Michael Ward and Ehsan Moharreri for their efforts working on this project in earlier stages. This research has been supported by the National Institutes of Health through grant number R35-GM119762 (to E.R.M.). This work used the Extreme Science and Engineering Discovery Environment (XSEDE) for HPC resources at the Texas Advanced Computing Center (TACC) through allocation TG-MCB140016. Additional computational resources for this work have been provided through the University of Connecticut Storrs HPC center.

