## Supplementary Information for "Atomistic Dynamics of a Viral Infection Process: Release of Membrane Lytic Peptides from a Non-Enveloped Virus"

Asis K. Jana, Eric R. May
Department of Molecular and Cell Biology, University of Connecticut, Storrs, CT 06269

### Supporting Information

Supporting information contains seven supporting figures (S1-S7) and four supporting movies (S1-S4).

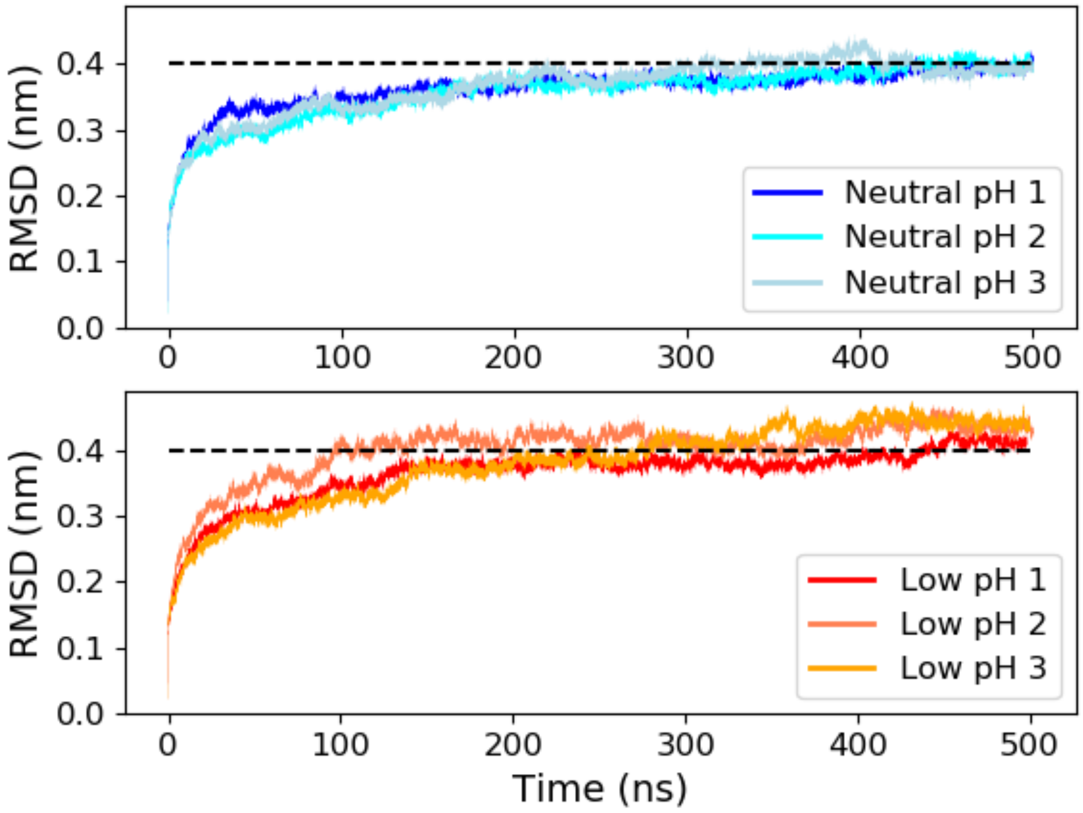


**Figure S1. RMSD of FHV pentameric capsid system.** RMSD was measured over three equilibrium simulation trajectories at each pH. The dotted line is drawn at 0.4 nm for a visual reference.

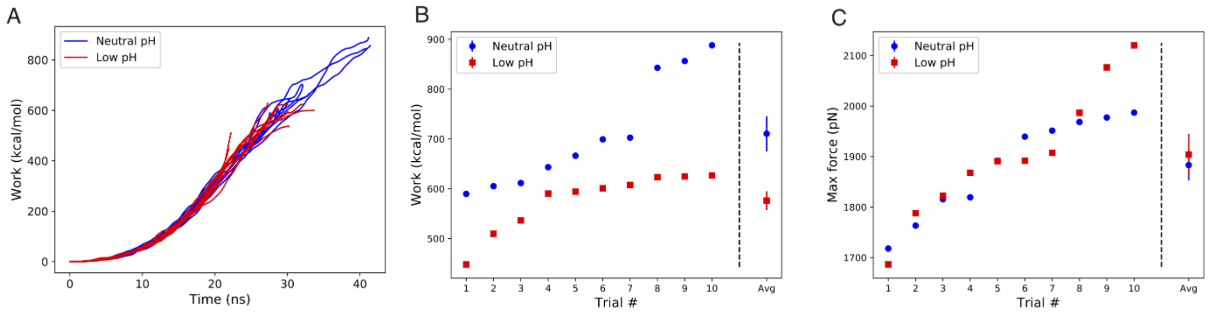


**Figure S2. SMD results using *coupled CV*.** A) The work versus time for the 10 trials at each pH. B) The final work values are sorted and presented from smallest to largest. The average over the trials is shown at the far right. C) The maximum force values are presented and ordered from smallest to largest. The average over the trials is shown at the far right. In (B-C) the error bars on the average values are the standard error of the means.

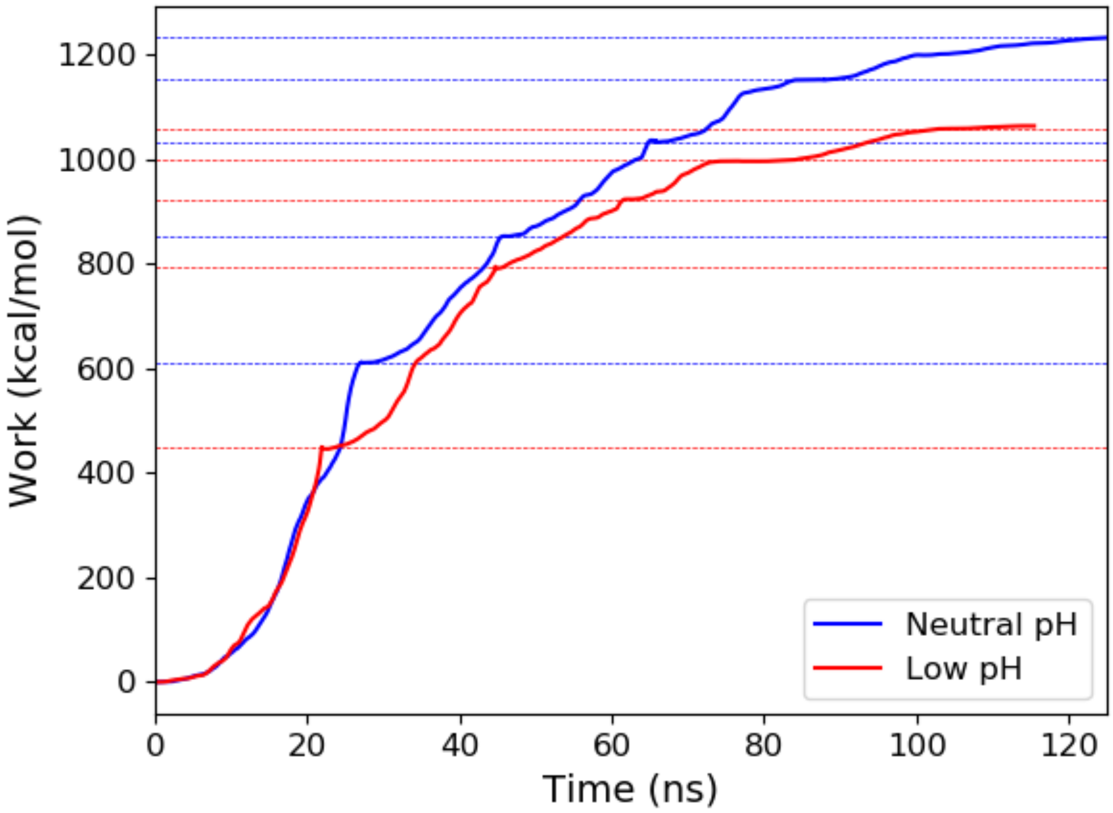


**Figure S3**. **SMD work required for externalization of five peptides.** Using the coupled CV, peptides were externalized from the capsid. Once a peptide was free from the capsid it was deleted, and the CV was redefined based on the remaining peptides and further biased. The dashed lines are drawn at the location of each peptide externalization event with blue corresponding to neutral pH and red corresponding to low pH.

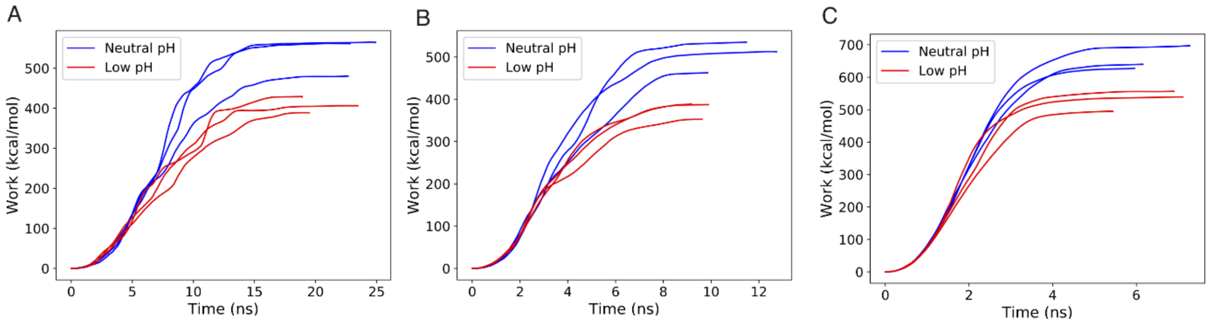


**Figure S4. SMD work using *single peptide CV* at increasing pulling rates.** Pulling rates are A) 0.2 nm/ns, B) 0.4 nm/ns and C) 0.8 nm/ns.

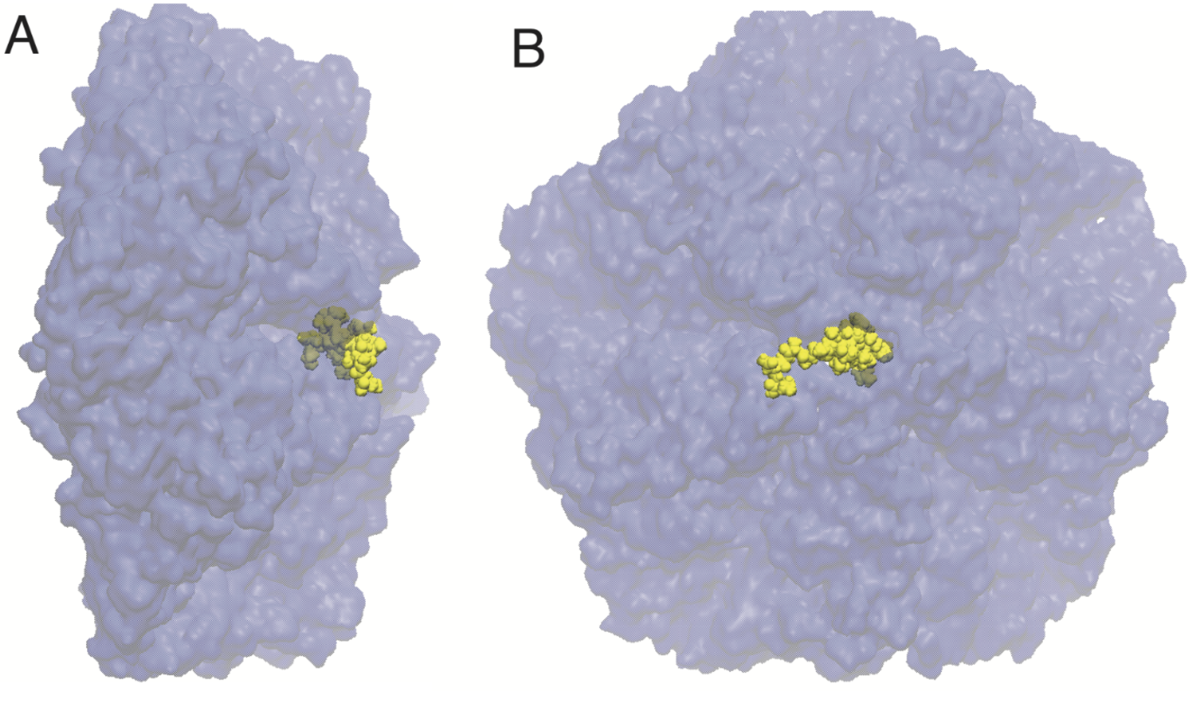


**Figure S5. Metastable state at low pH.** The final snapshot from umbrella window 35, which corresponds to CV=3.4 nm is shown from side (A) and top down (B) views. The capsid is depicted as a light blue surface and the externalizing γ peptide is depicted in yellow color using vdW representation.

```
CLUSTAL O(1.2.4) multiple sequence alignment

SP|P12870|CAPSD_FHV            MVNNNR------PRRQRAQRVVVTTTQTAPVPQQNVPRNGRRRRNRTRR----NRRRVRG 50
SP|Q9J7Z0|CAPSD_PAV            MVSRTK------NRRNKARKVVSRSTALVPMAPASQRTGP--------APRKPRKRN--- 43
SP|P04329|CAPSD_BBV            MVRNNN------RRRQRTQRIVTTTTQTAPVPQQNVPKQPRRRRNRARR----NRRQGRA 50
SP|P12871|CAPSD_NODAM          MVSKAARRRRAAPRQQQ--RQ----------QSNRASNQPRRRRA-----RR-TRRQQRM 42
SP|P12869|CAPSD_BOOLV          MT----------PRRQ--QRPKG---QLAKA---KQAKQP------LARSRRPRRRRRAA 36
TR|A0A140HER8|A0A140HER8_9VIRU MANNNQ------PKPRRQRRR-------NNAPKQQQNRAPRRRRNARRN----NR----- 38
TR|D0U4A0|D0U4A0_FHV           MVNNIK------PKRQRSQRVVVTTTQTAPIPQQNVPRNGRRRRNRARR----NRRRGRG 50
TR|D6PUS1|D6PUS1_9VIRU         MQRRQP----PRPRPQRTARVPS---QPVVALA---AAPPRRRRQQPRRPRRLRRQRRAV 50
                               *            : .   :                                  :

SP|P12870|CAPSD_FHV            MNMAALTRLSQPGLAFLKCAFAPPDFNTDPGKGIPDRFEGKVVSRKDVLNQSI-----SF 105
SP|Q9J7Z0|CAPSD_PAV            QALVRNPRLTDAGLAFLKCAFAAPDFSVDPGKGIPDNFHGRTLAIKDCNTTSV-----VF 98
SP|P04329|CAPSD_BBV            MNMGALTRLSQPGLAFLKCAFAPPDFNTDPGKGIPDRFEGKVVTRKDVLNQSI-----NF 105
SP|P12871|CAPSD_NODAM          AATNNMLKMSAPGLDFLKCAFASPDFSTDPGKGIPDKFQGLVLPKKHCLTQSI-----TF 97
SP|P12869|CAPSD_BOOLV          ITQNNLMMLSEPGLSFLKCAFASPDSNTDPGKGIPDNFEGKVLSQKNVYTETGVNFSGAT 96
TR|A0A140HER8|A0A140HER8_9VIRU M-ASSYMRLSEPGLAFLKCAFAPPDFNTDPGKGIPDRFEGKVVSRKDVLTAAT-----TF 92
TR|D0U4A0|D0U4A0_FHV           MNMAALSRLSQPGLAFLKCAFAPPDFQVRFGKCIPDRFEGKVVSRKDVLNQSI-----SF 105
TR|D6PUS1|D6PUS1_9VIRU         INTTALAQLTEPGLAFLKCAFAPPDFNTDPGKGIPDKFEGKVLSRKDVLTDTP-----SF 105
                                       ::  ** ******* ** ..  ** ***.*.* .:  *.  . :

SP|P12870|CAPSD_FHV            TAGQDTFILIAPTPGVAYWSASVPAGT-----FPTSATTFNPVNYPGFTSMFGTTSTSR- 159
SP|Q9J7Z0|CAPSD_PAV            TPNTDTYIVVAPVPGFAYFRAEVAVGA--------QPTTFVGVPYPTYATNFGAGSQNGL 150
SP|P04329|CAPSD_BBV            TANRDTFILIAPTPGVAYWVADVPAGT-----FPISTTTFNAVNFPGFNSMFGNAAASR- 159
SP|P12871|CAPSD_NODAM          TPGKQTMLLVAPIPGIACLKAEANVGAS------FSGVPLASVEFPGFDQLFGTSATDT- 150
SP|P12869|CAPSD_BOOLV          TQNVDTYIIVLPTPGVAFWRCIKTATAPAQPAALTTTDVFTAVPFPDFTSLFGTTATNR- 155
TR|A0A140HER8|A0A140HER8_9VIRU TAAKDTFILVAPTPGVAYWTAEVDSGT-----YPTSSTTFTAVVYPGFNSMFGETASAR- 146
TR|D0U4A0|D0U4A0_FHV           ESGKDTFILIAPTPGVVIWSASVDAGS-----FLTSTNTVRSSNYSGFTSMFGTTATSR- 159
TR|D6PUS1|D6PUS1_9VIRU         PANTDVYLLVLPTPGVAYWRLQKAAGT-----PIDTSDNFTAVLYPGFTSLFGTTAQSR- 159
                                   :. ::: * **..         :            .    :  :   **  :

SP|P12870|CAPSD_FHV            --SDQVSSFRYASMNVGIYPTSNLMQFAGSITVWKCPVKLSTVQFPVATD-PAT-----S 211
SP|Q9J7Z0|CAPSD_PAV            PAVNNYSKFRYASMACGLYPTSNMMQFSGSVQVWRVDLNLSEAVNPAVTAITPAPGVFAN 210
SP|P04329|CAPSD_BBV            --SDQVSSFRYASMNVGIYPTSNLMQFAGSITVWKCPVKLSNVQFPVATT-PAT-----S 211
SP|P12871|CAPSD_NODAM          --AANVTAFRYASMAAGVYPTSNLMQFAGSIQVYKIPLKQVLNSYSQTVATVPPTN-L-A 206
SP|P12869|CAPSD_BOOLV          --ADQVAAFRYASMNFGLYPTCNSTQYNGGISVWKGAVQMSTTQYPLDTT-PES-----S 207
TR|A0A140HER8|A0A140HER8_9VIRU --STNVSSFRYASMNVGIYPTSNMMQFGGSITVWKAPIQLSTNQFPIGADETAT-----S 199
TR|D0U4A0|D0U4A0_FHV           --SDQVSSFRYASMNVGIYPTSKLMQFTGSITVWKCPIKLSTVQFPVATE-PAT-----S 211
TR|D6PUS1|D6PUS1_9VIRU         --SANVSSFRYASMNCGLYPSSNMMQYGGSVSVWKVPIQMSTVQYPVSNS-VPT-----S 211
                                   : : ******  *:**:.:  *: *.: *::  ::

SP|P12870|CAPSD_FHV            SLVHTLVGLDGVLAV-GPDNFSESFIKGVFSQSACNEPDFEFNDILEGIQTLPPANVSLG 270
SP|Q9J7Z0|CAPSD_PAV            FVDKRINGLRGIRPLAPRDNYSGNFIDGAYTFAFDKSTDFEWCDFVRSLEFSE-SNVLGA 269
SP|P04329|CAPSD_BBV            ALVHTLVGLDGVLAV-GPDNFSESFIKGVFSQSVCNEPDFEFSDILEGIQTLPPANVTVA 270
SP|P12871|CAPSD_NODAM          QNTIAIDGLEALDAL-PNNNYSGSFIEGCYSQSVCNEPEFEFHPIMEGYASVPPANVTNA 265
SP|P12869|CAPSD_BOOLV          QLVHAITGLESALKV-GDENYSESFIDGVFTQSINGNAEFPFYPILEGVQTLPGQNVTVA 266
TR|A0A140HER8|A0A140HER8_9VIRU QLVHALQGLESVQAV-GSDNYSESFIKGVFSQSVCNEPEFEFKNIIEGVQTLPPQNVSLA 258
TR|D0U4A0|D0U4A0_FHV           SLVHILAGLDGVLAV-GPDNFSESFIKGVFSQSACNEPDFEFKDILEGIQTLPPSNVDIG 270
TR|D6PUS1|D6PUS1_9VIRU         QHSHALQGLEGVTFV-SQDNYTESFIKGMYASSVCNEPEFEFNNILEGVQTLPPVNVSIS 270
                                    : ** .   :   :*:: .**.* :: :   . :* :  ::..       **  .

SP|P12870|CAPSD_FHV            STGQPFTMDSGAEATSGVVGWGNMDTIVIRVSAPEGAVNSAILKAWSCIEYRPNPNAMLY 330
SP|Q9J7Z0|CAPSD_PAV            ATA-MKLLAPGGGTDTTLTGLGNVNTLVYKISTPTGAVNTAILRTWNCIELQPYTDSALF 328
SP|P04329|CAPSD_BBV            TSGQPFNLAAGAEAVSGIVGWGNMDTIVIRVSAPTGAVNSAILKTWACLEYRPNPNAMLY 330
SP|P12871|CAPSD_NODAM          QASMFTNLTF---SGARYTGLGDMDAIAILVTTPTGAVNTAVLKVWACVEYRPNPNSTLY 322
SP|P12869|CAPSD_BOOLV          QAGMPFSLDAGAATVAGFTGIGGMDAIFIKVTAAAGSVNTATIKTWACIEYRPNTNTALY 326
TR|A0A140HER8|A0A140HER8_9VIRU QSGQPFTFDAGSVSAAGIVGWGNMDTIIIRVSTAASAVNTAVVKTWACIEYRPNPNSAFY 318
TR|D0U4A0|D0U4A0_FHV           STGQPFTLSAGDENTSGTVGWGNMDTIVIRVSAPTGAVNSAILKAWSCIEYRPNPNAMLY 330
TR|D6PUS1|D6PUS1_9VIRU         QSGQPFGIDAGAENVCGITGFGNMDAIIIKVTTPTGATNVATFKNWACIEYRPTPNATLY 330
                                :.    :          .* *.::::   :::  .:.* * .: * *:* :*  :: ::

SP|P12870|CAPSD_FHV            QFGHDSPPLDEVALQEYRTVARSLPVAVIAAQNASMWERVKSIIKSSLAAASNIPGPIGV 390
SP|Q9J7Z0|CAPSD_PAV            QFSGVSPPFDPLALECYHNLKMRFPVAVSSRENSKFWEGVLRVLNQISGTLSVIPGPVGT 388
SP|P04329|CAPSD_BBV            QFGHDSPPCDEVALQEYRTVARSLPVAVIAAQNASMWERVKSIIKSSLAMASNVPGPIGI 390
SP|P12871|CAPSD_NODAM          EFARESPANDEYALAAYRKIARDIPIAVACKDNATFWERVRSILKSGLNFASTIPGPVGV 382
SP|P12869|CAPSD_BOOLV          KYAHDSPAEDIIALQQYRKVYKSLPVAVRAKLNANMWERVKRLLKAGLVAASYVPGPVGG 386
TR|A0A140HER8|A0A140HER8_9VIRU QFGHDSPRKDEYALEEYRRIAKQLPCAVEAAKNATMWERVKQIIKTGLSIASAVPGPIGM 378
TR|D0U4A0|D0U4A0_FHV           QFGHDSPPLDEVALQEYRTVAKSLPVAVIAAQNASMWERVKAIIKSSLAAASNIPGPIGV 390
TR|D6PUS1|D6PUS1_9VIRU         QYAHDSPPLDEIALREYRRVARCLPVAVPCAQNATMWERVKALLKSGLSALSAVPGPVGM 390
                               ::.  **  *  **  *: :   :* ** .  *:.:** *  :::      * :***:*

SP|P12870|CAPSD_FHV            AASGISGLSALFEGFGF 407
SP|Q9J7Z0|CAPSD_PAV            ISAGVHQLTGMYM---- 401
SP|P04329|CAPSD_BBV            AASGLSGLSALFEGFGF 407
SP|P12871|CAPSD_NODAM          AATGIKGIIETIGSLWV 399
SP|P12869|CAPSD_BOOLV          IATGVQHIGDLIAELSF 403
TR|A0A140HER8|A0A140HER8_9VIRU AATGLQGLSGLFTNLVI 395
TR|D0U4A0|D0U4A0_FHV           AASGISGLSALFEDFGF 407
TR|D6PUS1|D6PUS1_9VIRU         TASGIQGVSELLSGLFL 407
                                ::*:  :
```

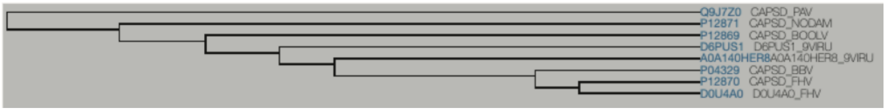


**Figure S6. Multiple sequence alignment of coat proteins from *alphanodavirus* taxa of *Nodavirus* family.** Histidine positions (215, 334) are highlighted in orange boxes. Below the MSA is the sequence tree. It can be seen that the only sequences which do not have histidine at same position as FHV (PAV and NODAM) are sequentially farthest from FHV. Abbreviations: FHV=Flock House Virus; PAV = Pariacoto virus; BBV=Black beetle virus; NODAM=Nodamura virus; BOOLV=Boolarra virus; A0A140HER8=Newington virus; D0U4A0=Drosophila melanogaster American nodavirus; D6PUS1=Alphanodavirus HB-2007.

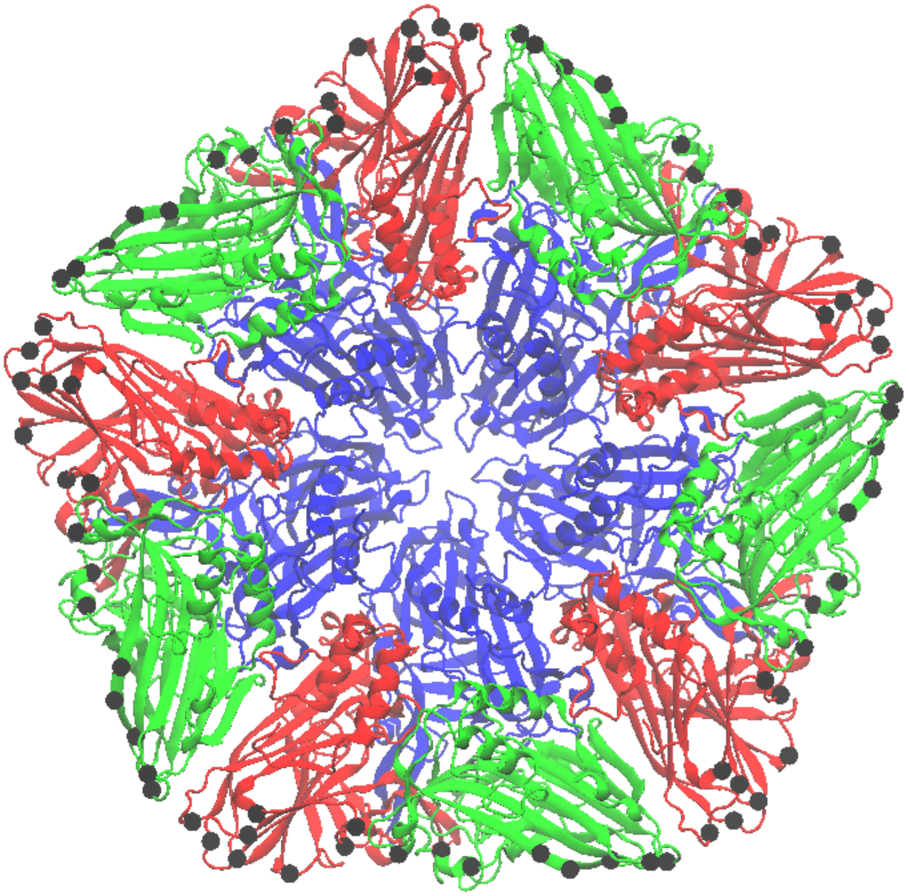

**Figure S7. Positional Restraints.** The following $C_\alpha$ atoms were restrained during all simulations to maintain the capsid curvature and are depicted by the black spheres. Chain B: residues 131, 183, 186, 225, 228, 232, 234, 304 and 307. Chain C: residues 100, 102, 147, 151, 180, 183, 310 and 328. Subunit coloring: chain A=blue, chain B=red and chain C=green.

**Movie S1. Equilibrium simulation of FHV pentamer at neutral pH for 500 ns.**

**Movie S2. Equilibrium simulation of FHV pentamer at low pH for 500 ns.**

**Movie S3. SMD trial for *coupled CV* at low pH.**

**Movie S4. SMD trial for *single peptide CV* at low pH.**
